# A chromosomal-level reference genome of the widely utilized *Coccidioides posadasii* laboratory strain “Silveira”

**DOI:** 10.1101/2021.05.19.444813

**Authors:** Marcus de Melo Teixeira, Jason E. Stajich, Jason W. Sahl, George R. Thompson, Austin V. Blackmon, Heather L. Mead, Paul Keim, Bridget M. Barker

## Abstract

Coccidioidomycosis is a common fungal disease that is endemic to arid and semi-arid regions of both American continents. *Coccidioides immitis* and *C. posadasii* are the etiological agents of the disease, also known as Valley Fever. For several decades, the *C. posadasii* strain Silveira has been used widely in vaccine studies, is the source strain for production of diagnostic antigens, and is a widely used experimental strain for functional studies. In 2009, the genome was sequenced using Sanger sequencing technology, and a draft assembly and annotation was made available. In the current study, the genome of the Silveira strain was sequenced using Single Molecule Real Time Sequencing (SMRT) PacBio technology, assembled into chromosomal-level contigs, genotyped, and the genome was reannotated using sophisticated and curated *in silico* tools. This high-quality genome sequencing effort has improved our understanding of chromosomal structure, gene set annotation, and lays the groundwork for identification of structural variants (e.g. transversions, translocations, and copy number variants), assessment of gene gain and loss, and comparison of transposable elements in future phylogenetic and population genomics studies.

## INTRODUCTION

Coccidioidomycosis is a multi-symptomatic mycotic disease affecting humans and other animals in arid and semi-arid regions in the Americas (Van Dyke *et al*. 2019). When aerosolized, conidia can be inhaled into the lungs of a susceptible host. In humans, the infection is asymptomatically controlled in 60% of infections (Nguyen *et al*. 2013). However if not controlled, the infection can progress from a mild pneumonia to a severe acute or chronic pulmonary infection. Moreover, the disease can disseminate into multiple organs (e.g. bones, skin, spleen, etc.) including the meninges, which is often fatal without treatment, and with the potential necessity of lifelong antifungal therapy (Galgiani *et al*. 2020). *Coccidioides immitis* and *C. posadasii* are the causative agents of coccidioidomycosis, and the two species diverged around 5.1 MYA (Engelthaler *et al*. 2016). Members of the *Coccidioides* genus are able to successfully interbreed, as evidenced by hybrids and patterns of introgression (Neafsey *et al*. 2010; Maxwell *et al*. 2019). Population structures within both species have been defined in several studies and generally reveal biogeographic patterns of distribution (Fisher *et al*. 2001; Teixeira *et al*. 2019). *C. immitis* is composed of up to three populations: San Joaquin (Central) Valley, San Diego/Mexico, and Washington, while three populations have been described within *C. posadasii*: Arizona, Texas/Mexico/South America and Caribbean (Teixeira *et al*. 2019).

The *C. posadasii* Silveira strain was collected from a patient of Dr. Charles E. Smith in 1951 (Friedman *et al*. 1955; Friedman *et al*. 1956). The patient had severe primary coccidioidomycosis with erythema nodosum, but recovered after several months of illness. Extensive testing in mice revealed that the strain was highly infective via the intraperitoneal route and caused nearly 100% lethal infections with as few as 100 conidia by 90 days (Friedman *et al*. 1956). Early studies analyzing the genetic diversity of *Coccidioides* via Restriction Fragment Length Polymorphisms (RFLPs) and Multi-Locus Sequencing Typing (MLST) of nuclear genes indicated that this strain grouped with the “non-California” population (Koufopanou *et al*. 1997). Although this strain was recovered from a patient residing in California at the time of diagnosis (prevalent area of *C. immitis*), subsequent genetic analysis demonstrated that this strain is *C. posadasii* (Fisher *et al*. 2002). Interestingly, the patient was a migrant farmworker that had previous travel history in Arizona in the year prior to diagnosis (chart review by GRT).

Despite the fact that this strain has been maintained in culture since the 1950s, many studies on virulence, vaccination challenge, and host response have been conducted without a loss of virulence in many labs. Mice challenged intranasally with 27 Silveira arthroconidia succumbed to infection rapidly, and the strain had an intermediate to high virulence when compared to other *C. immitis* and *C. posadasii* strains (Cox and Magee 2004). Silveira is able to initiate parasitic phase growth *in vitro* in Converse (Scalarone and Levine 1969) or RPMI media (Mead *et al*. 2019b), and early antigen preparations (coccidioidin and spherulin) for both intradermal cellular hypersensitivity and serologic tests were derived from this strain (Smith *et al*. 1961). Persistent skin reactivity was observed in people who recovered from primary coccidioidomycosis, which suggested that previous exposure to this fungus led to cellular immunity protection (Catanzaro *et al*. 1975). Early studies with this strain were seminal for correlating persistent protection to subsequent fungal exposure, and to the presence of viable *Coccidioides* cells in granulomas (Pappagianis *et al*. 1965; Pappagianis 1967). Later, Rebecca Cox *et al*. used this strain to demonstrate that hypersensitivity to coccidioidin in mice is mediated by cellular immunity and predicts protection against further infection (Cox *et al*. 1988).

Several immunization studies have been completed in various animal models using strain Silveira (Converse *et al*. 1962; Converse *et al*. 1964; Huppert *et al*. 1967; Peng *et al*. 1999; Hung *et al*.2000). An auxotrophic *C. posadasii* Silveira mutant generated by cobalt-60 irradiation was attenuated for virulence in murine models and showed protective capabilities. However, in the case of the auxotrophic mutant, virulence was restored over time (Pappagianis *et al*. 1961; Walch and Kalvoda 1971). Inactivated cells and proteins from Silveira have been used for vaccination also (Cox and Magee 2004). Mice immunized with formaldehyde-killed spherules, whole cell wall components, or proteins from Silveira showed a protective effect, even after challenge with different *Coccidioides* strains (Huppert *et al*.1967; Zimmermann *et al*. 1998; Peng *et al*. 1999). The nucleotide and protein sequences of Silveira have been further characterized and used for the production of recombinant proteins with the aim of developing an effective vaccine (Peng *et al*. 1999; Shubitz *et al*. 2002).

The Silveira strain was deposited at three fungal repositories. One was by M.S. Collins in the 1970s, from the lab of Dr. Pappagianis to the American Type Culture Collection (ATCC 28868) and the other from Dr. M.A Brandt (CDC) also originally from Dr. Pappagianis’ lab (but via Dr. R. Cox lab) to the Westerdijk Fungal Biodiversity Institute (CBS 113859). This strain is Lufenuron-resistant, and its growth is inhibited by polymyxin B and ambruticin (https://www.atcc.org/en/Products/All/28868.aspx). Presumably, each strain has been passaged both on plates and through mice multiple times over 70 years, but the exact accounting for each research group is unknown. Based on microsatellite profiles of strains from Dr. R. Cox and Dr. D. Pappagianis’ labs, there are 1-2 nucleotide length differences observed at 2 loci, K01 and K09 (Fisher *et al*. 2002). More recently, the strain was deposited by Dr. B.M. Barker to BEI resources (NR-48944), which was received from the lab of Dr. J.N. Galgiani, initially a gift from Dr. H. Levine, and this is the strain we sequenced in the current study. Interestingly, this strain was genotyped as *C. posadasii*, ARIZONA population and is further diverged from published data at two additional loci: GA37 and K07 (Fisher *et al*. 2002; Teixeira and Barker 2016).

The genome of this strain was previously sequenced using both Sanger-capillary and Illumina technologies to a 5.2x coverage, which was assembled into a 27.58 Mbp genome containing 54 nuclear scaffolds and 3 mitochondrial scaffolds with 10,228 protein-coding genes annotated (Neafsey *et al*.2010). Recently, the proteins from lysates and filtrates of both filamentous and parasitic phases of Silveira were sequenced using a GeLC-MS/MS approach (Mitchell *et al*. 2018). The authors reported 9,024 peptides from 734 previously annotated proteins, with 103 novel proteins described.

Previous sequencing efforts resulted in an unclosed draft genome and draft annotation (Neafsey *et al*. 2010). We therefore finished and re-annotated the genome to produce an updated assembly and annotation resource. A combination of both long-read Single Molecule Real Time Sequencing (SMRT) PacBio and paired short-read Illumina MiSeq technology was used to produce a chromosomal-level high-quality assembly. Our assembly approach resulted in five complete chromosomes, and a complete mitochondrial genome. The new annotation pipeline utilized a combination of transcriptomic and proteomic evidence, and generated a reduced number of total annotated genes compared to the original draft annotation.

## MATERIAL AND METHODS

### Strains and public data

We sequenced the *Coccidioides posadasii* strain Silveira that we obtained from the J.N. Galgiani collection, who originally received the strain from H. Levine, and we deposited this strain at BEI Resources (NR-48944). This strain has been deposited by other researchers previously to ATCC and Westerdijk Institute/CBS-KNAW resources (ATCC 28868 and CBS 113859). The first Sanger-based genome assembly of *Coccidioides posadasii* strain Silveira was downloaded from the NCBI (https://www.ncbi.nlm.nih.gov/nuccore/294654294) for comparisons, and the DNA used for sequencing was from the J.N. Galgiani lab, albeit an earlier passage than ours. Additionally, the *Coccidioides immitis* strains RS and WA_211 genome data was retrieved from NCBI (https://www.ncbi.nlm.nih.gov/nuccore/AAEC00000000v3, https://www.ncbi.nlm.nih.gov/nuccore/1695747985). The annotation for RS was last updated in March 2015.

### DNA extraction

DNA extraction for short and long read sequencing was initiated by growing the Silveira strain from arthroconidia in liquid 2xGYE (2% glucose, 1% Difco Yeast Extract in dddH2O) for 120 hours and harvesting by centrifugation and washing twice with sterile 1xPBS. DNA for the short read library was obtained as previously described (Mead *et al*. 2019a). DNA for the long read library was obtained by freezing mycelia in liquid nitrogen and ground in a sterile mortar and pestle to a fine powder in a biological safety cabinet in a biosafety level three laboratory following the U’Ren high molecular weight (HMW) fungal DNA laboratory protocol (U’Ren and Moore 2018). Briefly, approximately three grams of ground fungal biomass was added to 14 mL SDS buffer (20mM Tris-HCl pH 7.5, 1mM EDTA pH 8, 0.5% w/v SDS) and incubated at 65°C, followed by addition of 0.5X potassium-acetate (5M KOAc, pH 7.5) to SDS buffer and incubation in ice for 30 minutes. The sample was centrifuged and the supernatant was subjected to isopropanol precipitation and ethanol cleanup. The resulting pellet was suspended in TE buffer (Tris-HCl 1mM, EDTA 0.5mM) and RNase treated, followed by phenol/chloroform extraction and ethanol precipitation. The final DNA pellet was slowly suspended in a minimal volume of low salt TE for sequencing. HMW DNA was quality checked for size using chromatin electrophoresis gel, for purity on Nanodrop 2000 (Thermo Fisher, USA) for 260/280 and 260/230 ratios, and quantified by Qubit (Invitrogen, USA) fluorometry.

### Sequencing

Short read sequencing was performed on a MiSeq (Illumina) instrument at The Translation Genomics Research Institute, Flagstaff, Arizona, as previously described (Teixeira *et al*. 2019). Long read sequencing was performed at the University of Arizona Genomics, Tucson, Arizona, core facility. Briefly, a sequencing library was constructed from 6ug of HMW DNA following the PacBio (Pacific Biosciences, USA) protocol for use with the SMRTbell Express Template Prep Kit. The ligated library templates were size selected on Sage BluePippin instrument (Sage Science, USA) for selection of fragments of 17 Kbp and larger, which is appropriate given the predicted smallest chromosome is greater than 4 Mbp (34). The final purified sequencing library yield was 980ng with a final mode size of 38.3 Kbp as determined by Fragment Analyzer (Agilent Technologies, USA). The final library was bound with PacBio polymerase and sequencing primer v3 using manufacturers’ methods. One 1M v2 SMRT cell was loaded with a bound library at a loading concentration of 7pmol/cell followed by a 10-hour sequencing run on the PacBio Sequel Instrument at the University of Arizona, Tucson, AZ, resulting in ~475x coverage. Additionally, ~100X sequencing coverage with Illumina short reads was generated on Illumina MiSeq (2 × 300 bp mode) instrument in a high output mode (Illumina, San Diego, CA) to support PacBio read correction and assembly polishing as needed.

### Assembly

The reference genome assembly pipeline was comprised of five steps: **1)** PacBio reads were corrected using Illumina reads and the tool LoRDEC v 0.9 (Salmela and Rivals 2014); **2)** Corrected PacBio reads were assembled with Canu software v 1.7.1 (Koren *et al*. 2017); **3)** An additional round of assembly using Illumina short reads combined with the Canu PacBio scaffolds as trusted contigs in the assembler SPAdes v 3.13 (Bankevich *et al*. 2012); **4)** five rounds of Pilon (v 1.24) corrections improved the genome assembly to further reduce nucleotide base error variants (Walker *et al*. 2014), and; **5)** we further scaffolded the assemblies to existing *Coccidioides immitis* strain RS reference genome to compare synteny (Sharpton *et al*. 2009) with RaGOO v 1.1 (Alonge *et al*. 2019). We evaluated and compared assembly accuracy with the hybrid assembly method MaSuRCA v 3.3.0 (Zimin *et al*. 2013; Zimin *et al*.2017), which incorporates the high performance long read assembler Flye v 2.5 and integrates short read correction of reads and assembly into the pipeline (Kolmogorov *et al*. 2019). Assembly quality was assessed by using summary statistics, the number of complete alignments of conserved fungal proteins using BUSCO v 2.0 (Simao *et al*. 2015; Vanderlinde *et al*. 2019), RNASeq read mapping from existing libraries (Whiston *et al*. 2012), and utilizing existing and *de novo* annotations of *Coccidioides* genomes (Sharpton *et al*. 2009; Neafsey *et al*. 2010). Transposable elements and low complexity DNA sequences were assessed using RepeatMasker v 4.1.1 (Smit 2020). Telomeric repeats were identified using the FindTelomers python script (https://github.com/JanaSperschneider/FindTelomeres). The quality of newly and previously *Coccidioides* assembled genomes were assessed using the QUAST v5 pipeline (Gurevich *et al*. 2013). The sequence composition of predicted 18S rRNA genes was performed with SSU-Align v 0.1.1 (eddylab.org/software/ssu-align/). The assembly of the mitochondrial DNA using Illumina reads was performed according to our protocol described previously (de Melo Teixeira *et al*.2021). Long read mapping to the mitochondrial scaffolds were accomplished using the Minimap2 algorithm (Li 2018) and read aliments were visualized using the Tablet software (Milne *et al*. 2012). Dot Plot comparisons were performed using Gepard (Krumsiek *et al*. 2007) and MAFFT v 7 (Katoh and Standley 2013).

### Annotation

The Funannotate pipeline v 1.7.4 (Stajich and Palmer 2020) was used to automate *ab initio* gene predictor training using BRAKER1 (Hoff *et al*. 2016), Augustus v 3.3 (Stanke *et al*. 2006), and GeneMark-ET v 4.57 (Ter-Hovhannisyan *et al*. 2008). This pipeline generated *de novo* assembly of transcripts with Trinity v 2.10 (Haas *et al*. 2013) to examine variation in exons used in isoforms, and aligned to the genome. This evidence was used as input data for EVidence Modeler (EVM) software v1.1.1 to generate a consensus set of predicted gene models (Haas *et al*. 2008). Gene models were filtered for length, spanning gaps and transposable elements using a *Coccidioides*-specific library of repetitive DNA to further clean the dataset (Kirkland *et al*. 2018). AntiSMASH 5.1.1 (fungismash) was used to identify biosynthetic gene clusters (Blin *et al*. 2019). The mitochondrial annotation was performed using the MFannot and RNAweasel pipelines available at https://megasun.bch.umontreal.ca/.

### Phylogenomic classification

To characterize the phylogenetic relationship of the Silveira strain, we first used the assembled genome as reference for Illumina read mapping and SNP variant calling utilizing the Northern Arizona SNP Pipeline (NASP) v 1.0 (Sahl *et al*. 2016). We mapped 61 available genomes of *C. posadasii* against our five Silveira scaffolds using the Burrows-Wheeler Aligner (v 0.7.7) tool (Li and Durbin 2009). SNPs were called using the GATK (v 3.3.0) toolkit (McKenna *et al*. 2010) using previous NASP protocols (SNP filtering and indel-based realignment) developed by our group for *Coccidioides* genotyping (Teixeira *et al*. 2019). A total of 258,470 SNPs were retrieved and submitted for unrooted phylogenetic analysis via maximum-likelihood method implemented in the IQTREE software v 1.6.12 (Nguyen *et al*. 2015). The best-fit model was set according to Bayesian Information Criterion to TN+F+ASC+R6 and the phylogenetic signal was tested using both Shimodaira–Hasegawa approximate likelihood ratio test (SH-aLRT) and ultrafast bootstrap support (Anisimova and Gascuel 2006; Minh *et al*. 2013). The phylogenetic tree was visualized using the Figtree software (http://tree.bio.ed.ac.uk/software/figtree/). The population structure of *C. posadasii* and the proportion of admixture of the Silveira strain was inferred based on unlinked SNPs using the fastSTRUCTURE software (Raj *et al*. 2014). The best population scenario (or *K*) was calculated using the choosek.py application based on lowest marginal likelihood and cross-validation test. The proportion of the admixture of each individual was plotted using the Structure Plot v2.0 pipeline (Ramasamy *et al*. 2014).

### Data availability

Sequence data and the final assembly were submitted to GenBank under BioProject PRJNA664774. PacBio Sequel sequencing data are found at SRR 9644375. Illumina MiSeq sequencing data are found at SRR 9644374. Final nuclear assembly is deposited in accession CP075068-CP075075 and the MT genome as accession (TBD).

## RESULTS AND DISCUSSION

### Sequencing and Assembly

PacBio Sequel sequencing yielded ~10.6 Gbp of raw data from over 643,000 reads with an average read length (N50) of ~25 Kbp. Illumina MiSeq sequencing yielded ~2.8 Gbp of raw data and >12 million paired-end short reads. We used both short-read Illumina and long-read PacBio data to assemble the genome using short-read-corrected (Pilon), long read (Canu), and a long-read alone (Flye) assembly approaches. The genome was assembled with Canu on the corrected PacBio sequencing data into 9 scaffolds totaling 28.27 Mbp; scaffolds 1-5 representing chromosomes 1-5, scaffold 6 as the mtDNA, and scaffolds 7-9 are likely supernumerary chromosomes that we named chromosomes 6-8 (Table 1). The L50 metric for this assembly is 2 Mb, the N50 is 8.07 and the longest scaffold is 8.34 Mbp. A search for telomeres using Flye identified 10 telomeric repeats on chromosomes 1-5. Sequence searches for 18S rRNA genes identified five loci, two of them were complete with an additional that was nearly complete. By comparing the assembly metrics of the Silveira strain between Sanger and PacBio/Illumina assemblies, we observed a complete genome structure of this fungus consistent with its known chromosomal composition (Table 2, Supplemental Figure 1) (Pan and Cole 1992). The genome size of the assembly using Canu is 690 Kbp longer than the previous Sanger assembly, which is contained in 54 nuclear genome scaffolds, with three mitochondrial scaffolds. The Canu assembly produced five large scaffolds which have nine telomeric repeats (TTAGGG/CCCTAA)_n_ at both ends of scaffolds 1, 2, 3 and 5. On scaffold 4 this tandem repeat was only found at the forward strand in the Canu assembly (Figure 1D). According to the densitometry analysis of Pulsed-Field Gel Electrophoresis (PFGE), the *C. posadasii* Silveira should have 4 chromosomal bands (Pan and Cole 1992). However, because scaffolds 1 and 2 are 8.34 Mbp and 8.07 Mbp, respectively, these minor chromosomal size differences would be difficult to discriminate using PGFE.

**Table 1.**
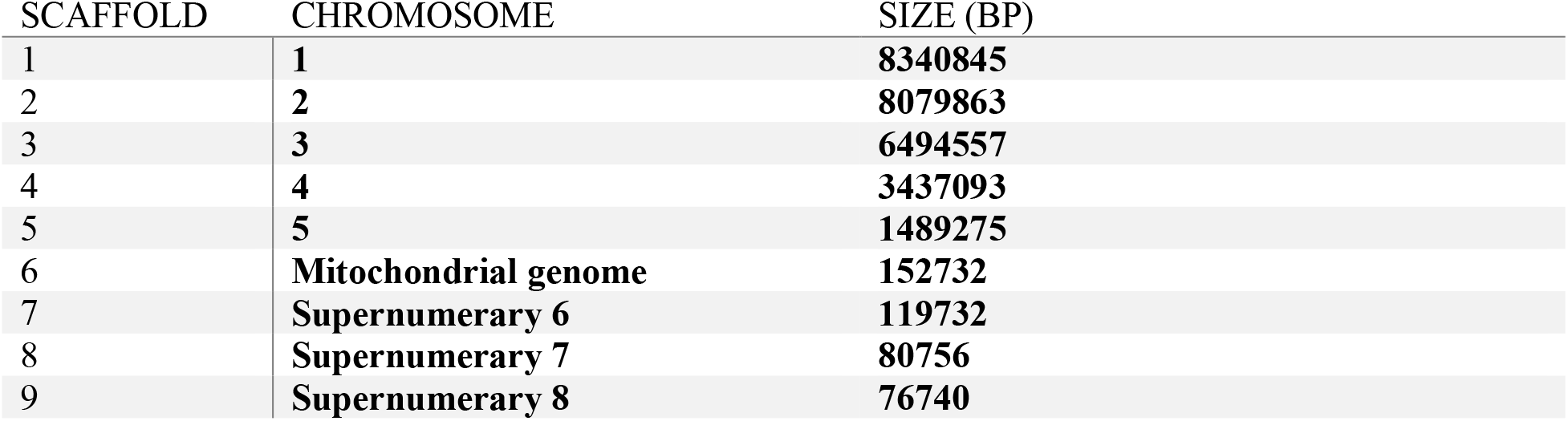
Chromosome Sizes.

**Table 2.** Comparison of Silveira Genomes.

|  | Silveira Sanger<br>2007 | Silveira CANU<br>2020 |
| --- | --- | --- |
| **Contigs** | 57 | 9 |
| **GC%** | 46.34% | 46.44% |
| **Predicted**<br>**Protein-**<br>**coding Genes** | 10,228 | 8,491 |
| **Median Gene**<br>**Length (bp)** | 2109 | 1810.5 |
| **Total number**<br>**of introns** | 20607 | 17004 |
| **Total # of**<br>**Exons** | 32511 | 26348 |
| **Total # of**<br>**CDS** | 30840 | 26216 |
| **tRNAs** | **119** | **111** |

**Figure 1.**
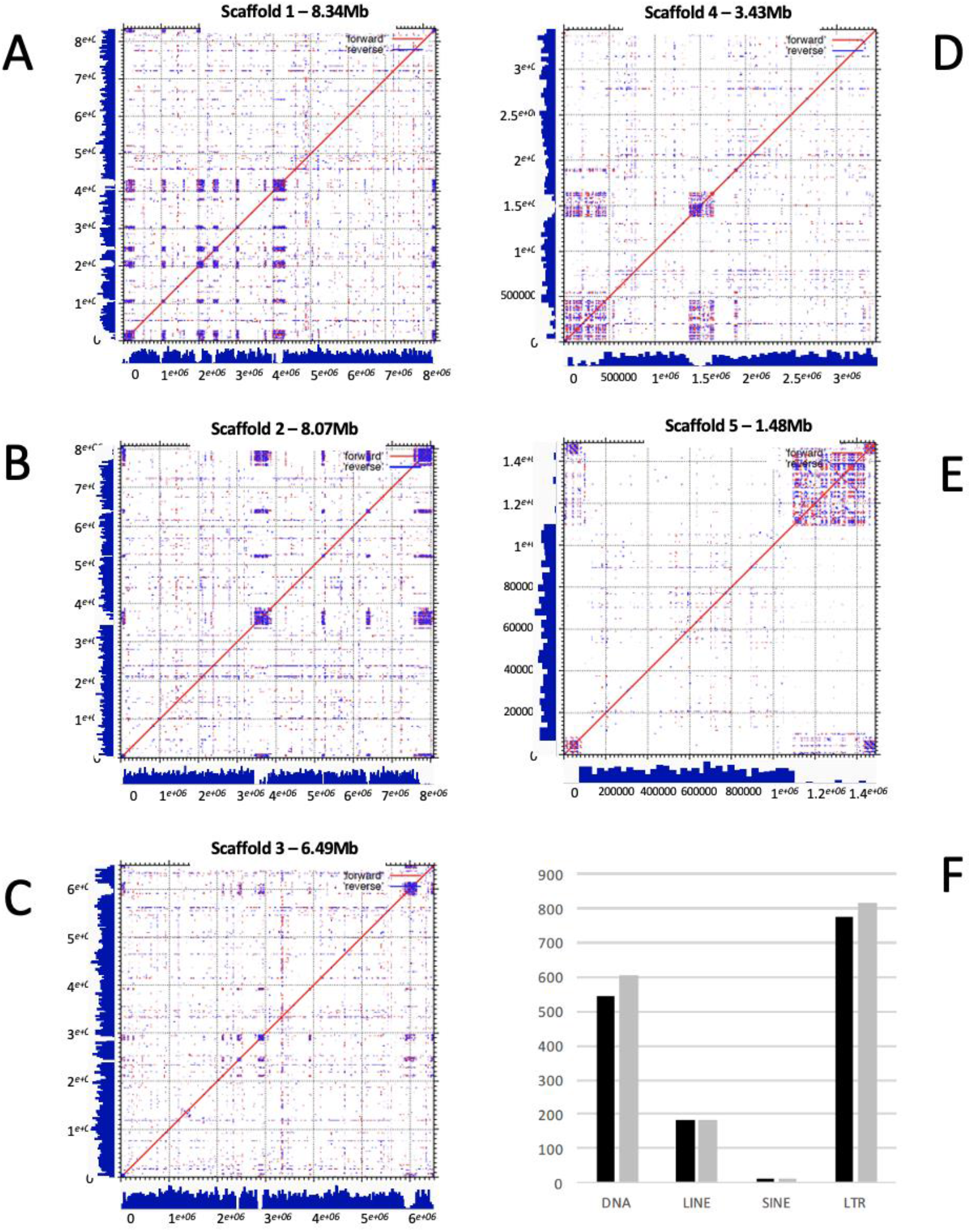
The gene density and repeats for Canu assemblies. Blue histograms on X/Y axes represent gene densitities. Forward and reverse similarity blocks are indicated by red and blue dots, respectively. A central high repeated sequence regions is marked as a putative centromere. Canonical telomeric repeats are indicated at both 5’ and 3’ ends. **A) Dot Plot of Chromosome I. B**) **Dot Plot of Chromosome II. C) Dot Plot of Chromosome III. D) Dot Plot of Chromosome IV. E) Dot Plot of Chromosome V. F) Classes of repeats.**

### Repetitive DNA

Highly-repeated DNA sequences play a pivotal role in genome architecture, evolution and adaptation by modulating gene activity, and chromosomal rearrangements (Castanera *et al*. 2016; Sinha *et al*. 2017; Nieuwenhuis *et al*. 2019). The newly assembled Silveira genome has 17.12 % total repetitive DNA assessed with RepeatMasker, while the previous version had 15.91%. The new assembly has 1,614 copies of various transposable elements (TEs), whereas the earlier version has 1,511. The most impacted TE classes were LTR retrotransposons and DNA transposons (Figure 1F). This difference is likely due to improvements in the assembly because long read technology can cover long stretches of repetitive DNA that may have been collapsed during the assembly process of Sanger and Illumina short reads. There is strong evidence for sub-telomeric repeats, as demonstrated by the presence of low complexity sequences adjacent to terminal positions of the five larger scaffolds (Figure 1). We observed an accumulation of repetitive DNA and lack of protein-coding genes in central positions of the scaffolds, which may be centromeric repeats. We have found that rnd motifs were abundant in the putative centromeres; however, chromatin immunoprecipitation followed by deep sequencing are needed to determine whether those central repeats are in fact the centromeres of *C. posadasii*.

### Chromosome Structure

Although we used multiple approaches to assemble genomes, the assemblies produced were largely in agreement, creating five large contigs (chromosomes). The Flye algorithm assemblies generated marginally longer contigs. For all five chromosomes, there were possible internal centromeric locations and canonical telomeres at the end (Figure 1). Chromosome IV had flanking telomeres for the Flye assembly, but only on the 5’ end of the Canu contig. This discrepancy is likely due to an assembly error with Canu algorithm, and the Flye-assembled Chromosome IV is complete.

### C. immitis Genome Comparison

A comparison of the assembled Silveira genome to the published RS (*C. immitis*) genome suggests that RS has slightly larger genome, with note translocations and inversions (~2 Mb, Supplemental Figure 1C). This difference could be due to sequencing technology and assembly methods, but may in fact represent a true difference between species, because we see similar patterns when looking at another *C. immitis* genome WA_211 (Supplemental Figure 1 C-D). Variation in genome size between two divergent species is possible, as well as differences among isolates of the same species. Alternatively, Silveira has been a laboratory isolate since the 1950’s and has likely undergone micro-evolutionary changes due to *in vitro* selection for laboratory cultivation. Different isolates of the *Cryptococcus neoformans* var. *grubii* H99 genotype obtained from different laboratories display remarkable genetic variation related to microevolutionary processes of *in vitro* passage (Janbon *et al*. 2014). Genome reduction and other micro-evolutionary changes due to selective forces in the laboratory have been documented for other fungal pathogens (Chen *et al*. 2017; Ene *et al*. 2018). As high-fidelity long read technologies become increasing available, more accurate estimates of both genome size for *Coccidioides* spp. and genome variation among isolates will be possible.

While most of the RS and Silveira genome assembles are syntenic (Supplemental Figure 1), there is a notable exception for Silveira chromosome III (Figure 2). Chromosome III aligns to the RS genome, but spans two RS contigs. RS supercontig 3.1 contains the 5’ 30% of Silveira chromosome III (~3 Mb) with the remainder found on RS supercontig 3.2. The 3.2 supercontig alignment has a chromosomal rearrangement that is consistent with an inversion of ~3 Mb. Genome assembly algorithms have difficulties in processing direct and indirect repeated sequences, and this can result in erroneous results (Tørresen *et al*. 2019). However, in this particular case, the inversion junction points are not associated with highly repetitive sequence. We manually examined the Silveira reads and assemblies around the junction points, and we found clear evidence that our chromosome III assembly is correct. Similar raw read data is not available for the RS genome, and these data will be required to confirm this inversion. Chromosomal inversions and translocations can greatly influence meiotic processes by disrupting homolog pairing and recombination. Such reproductive barriers often exist between species and higher taxonomic relationships, limiting gene flow and allowing for differentiation and reinforcement of species boundaries. Again, the long *in vitro* culture history of Silveira leaves open the possibility that the apparent inversion may be a lab-specific or strain-specific genomic feature that will not be observed in all isolates of *C. posadasii*. Conversely, RS is also a lab strain that has been cultured under laboratory conditions since the 1950’s, with extensive evidence of hybridization and introgression, and thus the inversion could be specific for strain RS (Sharpton *et al*. 2009; Neafsey *et al*. 2010); however, we require additional long read sequencing and assemblies to answer this specific question.

**Figure 2.**
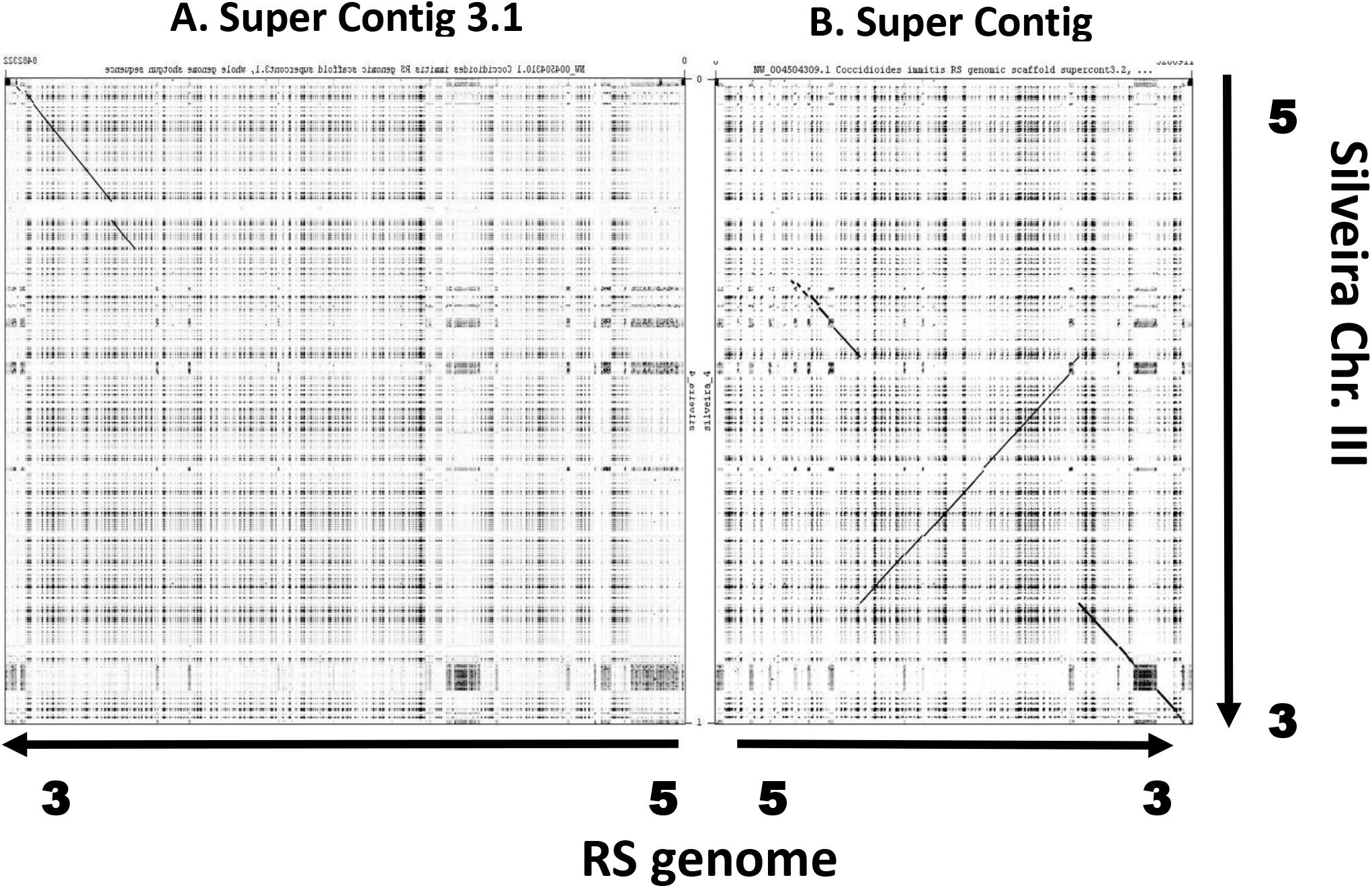
Silveira (*C. posadasii*) Chromosome III Dot Plot Similarity to the RS (*C. immitis*) Genome. The homologous regions of the Silveira chromosome III (6.5 mbp) are distributed across two super contigs of the RS genome assembly and not completely collinear. A) RS super contig 3.1 contains about 30% the 5’ Silveira chromosome III homologous sequences, with the remainder not homologous to chromosome III. Within the homologous region there a large gap of non-homology. B) RS super contig 3.2 contains about 70% of Silveira chromosome III homologous sequences including a highly repetitive region at the Silveira 3’ end. The arrangement of the cross species homologous regions are not completely collinear with an apparent inversion of about half.

### Mitochondrial Genome

Scaffold 6 corresponds to the mitochondrial genome, which was assembled to a size of 152 Kbp, which is almost double that of the Illumina assembly version that yields a 74 Kbp circular mapping genome (Figure 3) (de Melo Teixeira *et al*. 2021). These results indicate assembly inconsistencies between the two sequencing technologies. We mapped Pacbio reads against the PacBio mitogenome assembly, and we identified long reads that span the mitogenome of the Canu assembly (Supplemental Figure 2). A self-alignment of the 152 Kbp assembly reveals a dimeric and circular structure that could represent a biologically relevant molecule. We find the same 14 protein-coding mitochondrial genes that were annotated previously (de Melo Teixeira *et al*. 2021) as part of the ubiquinone oxidoreductase, cytochrome b, cytochrome oxidase and ATP synthase protein complexes. Small and large ribosomal RNAs (*rns* and *rnl*), the RNAseP subunit (rnpB), as well as 24 tRNAs were also annotated.

**Figure 3.**
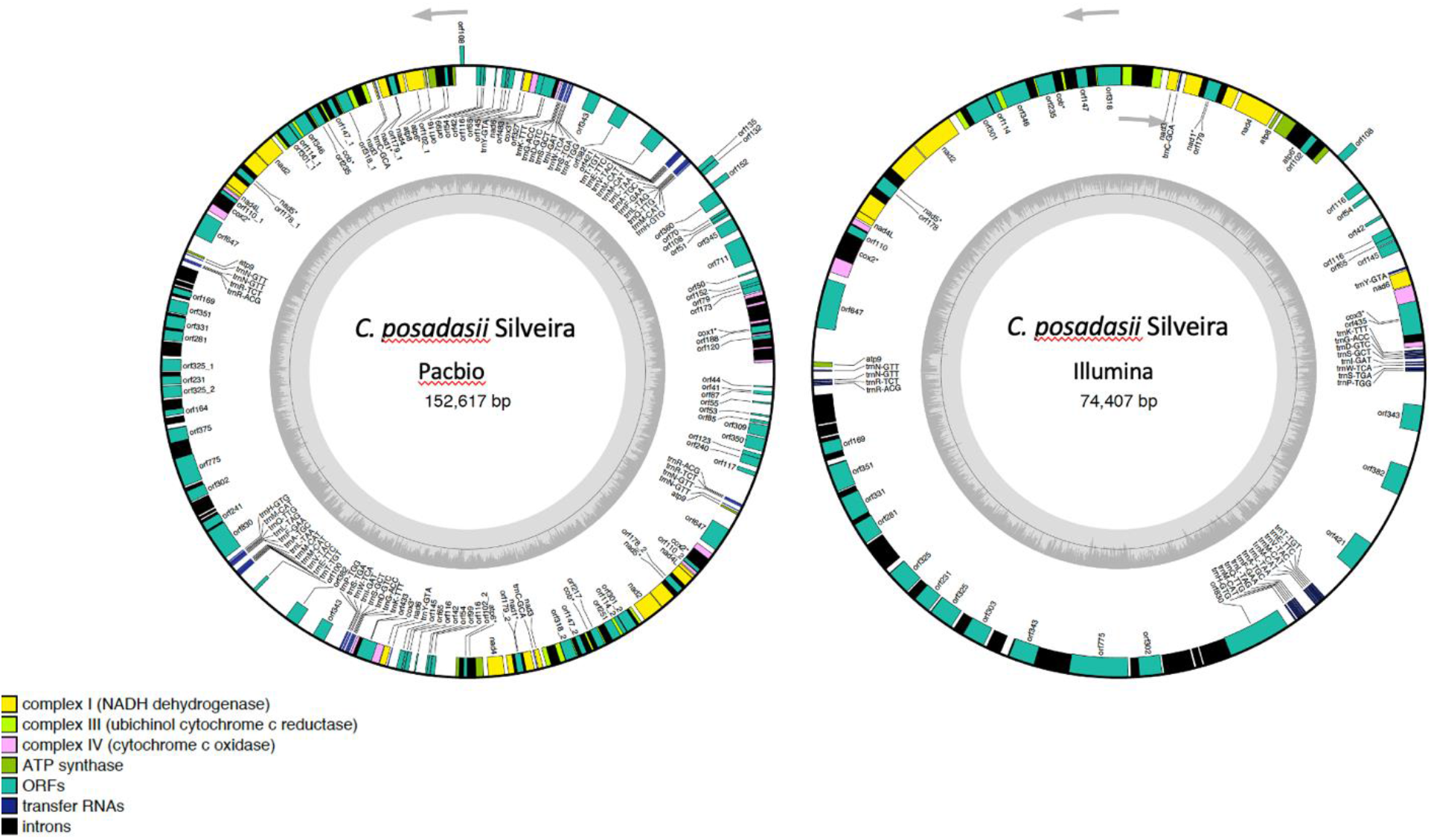
Circular mitochondrial genome assemblies of the *C. posadasii* Silveira strains using both Pacbio and Illumina technologies. The circular plots show the core genes as part of the ubiquinone oxidoreductase, cytochrome b, cytochrome oxidase and ATP synthase protein complexes. Small and large ribosomal RNAs (*rns* and *rnl*), the RNAseP subunit (rnpB), as well as tRNAs are also displayed. Additional ORFs and introns are showed along the circular map.

### Additional minor contigs

Lastly, three remaining scaffolds (7 to 9) are less than 119 Kbp total, but scaffolds 7 and 8 do contain coding sequences that were not found in the five chromosome or mitochondrial DNA assemblies. At least some of these small contigs contain real genes that failed to assemble into the larger chromosomes. Until additional sequences are completed from other isolates, we propose that these are supernumerary chromosomes (Coleman *et al*. 2009).

### Gene Models

Several metrics suggest a substantial improvement for the gene models of the Silveira genome. We have identified 8,491 total gene models in the new Silveira assembly, which is lower than the previous draft version of this genome, which had 10,228 gene models. The number of exons, mean exon length, and overlapping genes are lower in the new Silveira assembly than the previous one (Table 2). A more fragmented assembly would create discontinuous exons, and thus increase the number of gene models. We have found 207 genes with at least 1 isoform, suggesting alternative splicing. Additionally, the multiple training steps using mRNA sequence data and proteomics, along with the more sophisticated gene predictors implemented in the funannotate pipeline has increased the overall confidenc of gene models.

### Secondary Metabolism

Previous genomic analysis identified genes associated with secondary metabolite (SM) production that are shared between the two *Coccidioides* species, and have experienced positive selection (Sharpton *et al*. 2009). This observed evolutionary pattern might help the organism survive the harsh microenvironments in desert areas where the organism lives, or these secondary metabolites may be important for host-pathogen interactions (Perrin *et al*. 2007; Narra *et al*. 2016). Previous analysis suggested that *C. immitis* and *C. posadasii* have 22 and 21 SM clusters (Shang *et al*. 2016) although strain information was not given. We retrieved 23 SM gene clusters using antiSMASH analysis on our new Silveira genome assembly. The biosynthetic class polyketide synthase (PKS) is overrepresented in *Coccidioides* and these sequences display similarity with other well-known PKS clusters in the Ascomycota such as chaetoviridin E, depudecin, nidulanin A, cichorine, shanorellin, aflatoxin, stipitatic acid and leucino-statin A. The genomes of closely-related dermatophyte species also contain 23-25 SM clusters, but only 9 of these clusters are shared between *Coccidioides* and the dermatophytes, which might be related to the diverse ecological niches occupied by the two groups of Onygenalean fungi (Martinez *et al*. 2012).

### Phylogenomic characterization

Previous studies analyzing the genetic background of the Silveira strains based on microsatellite markers suggested that this strain belonged to the *C. posadasii* Arizona population (Teixeira and Barker 2016). With the advances of genome-based typing methods in *Coccidioides*, novel phylogenetic groups were defined within this species as follows: ARIZONA, Clade AZ1, Clade Col. Springs/GT162, TX/MX/SA and CARIBE (Teixeira *et al*. 2019). By adding the Silveira as a reference genome for whole genome typing we confirmed that this strain grouped within *C. posadasii*, ARIZONA clade based on the unrooted phylogenetic tree (Supp Figure 3A). By applying population genetics methods we have also observed that the Silveira strain belongs to the ARIZONA population and no admixture was found (Supplemental Figure 3B).

## SUMMARY

In the current study, the genome of the Silveira strain was sequenced using long read sequencing technology, and assembled into chromosomal-level contigs, and reannotated using current technology. This has dramatically improved our understanding of chromosomal structure, gene set annotation, and was a necessary step toward future genomics studies. Our assembly approach resulted in five complete chromosomes, a complete mitochondrial genome, and an indication of supernumerary chromosomes. The new annotation pipeline utilized a combination of transcriptomic and proteomic evidence, and generated a reduced number of total annotated genes compared to the original draft annotation. These critical advancements will allow for a better understanding of the evolution of this important endemic fungal pathogen.

## Acknowledgments

The authors wish to acknowledge Dr. Jana U’Ren for sharing High Molecular Weight DNA isolation protocol, Dr. Dave Kudrna (U Arizona) for advice and sequencing on PacBio. Funding to PK ABRC / ADHS18-198851. BMB was partly funded by R21AI28536-01AI. JES. is a CIFAR Fellow in the program Fungal Kingdom: Threats and Opportunities and was supported by University of California MRPI grants MRP-17-454959 “UC Valley Fever Research Initiative” and VFR-19-633952 “Investigating fundamental gaps in Valley Fever knowledge” and United States Department of Agriculture - National Institute of Food and Agriculture Hatch Project CA-R-PPA-5062-H.

---

**Supplementary figure 1.**
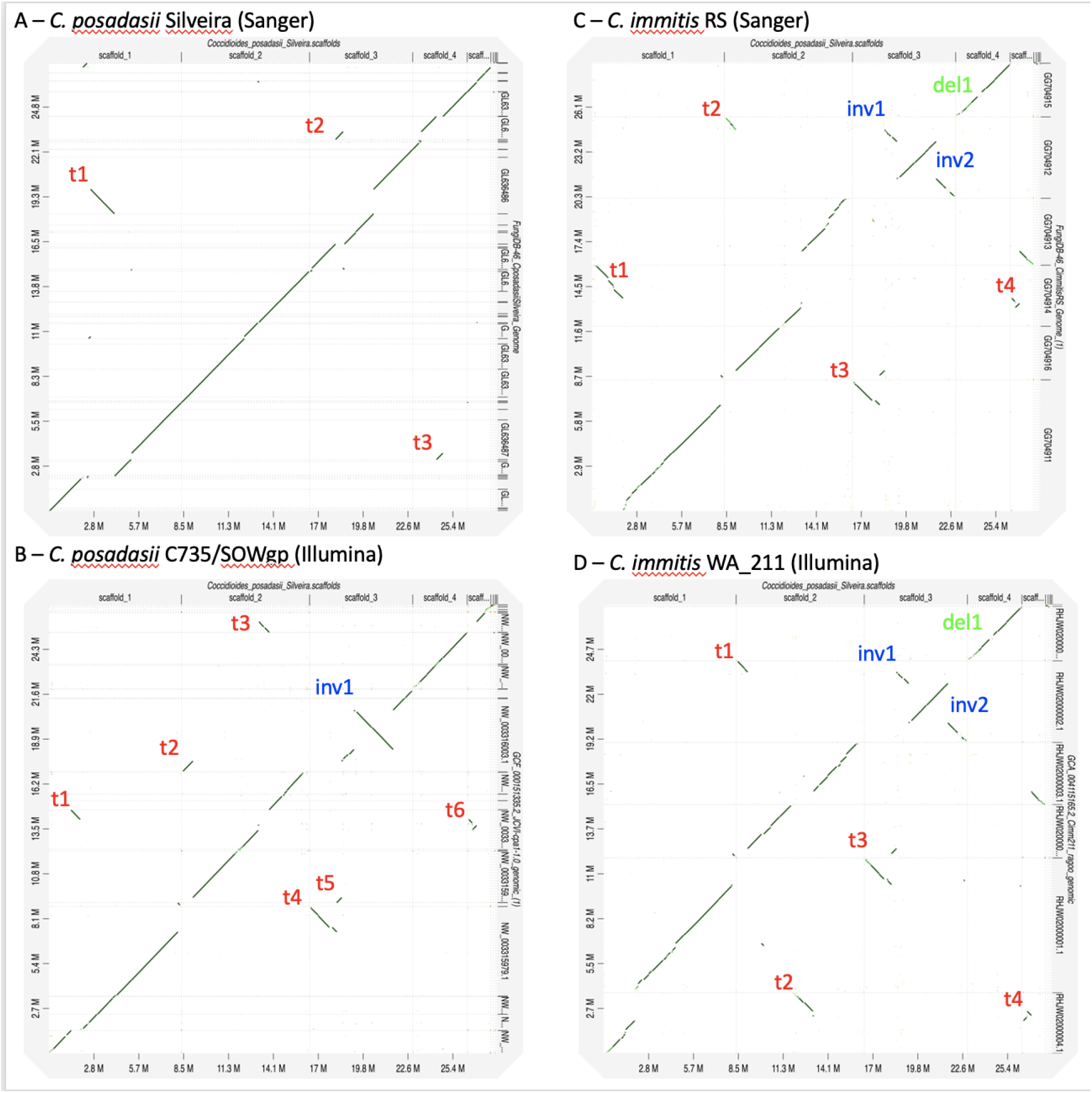
Whole-genome dot-plot alignments of the *C. posadasii* Silveira Pacbio assembly against: A) *C. posadasii* Silveira Sanger assembly, B) *C. posadasii* C735/SOWgp Illumina assembly, C) *C. immitis* RS Sanger assembly and D) *C. immitis* WA_211 Illumina assembly. The main chromosomal rearrangements such as inversions (inv), translocations (t) and deletions (del) are highlighted in different colors.

**Suppl Figure 2:**
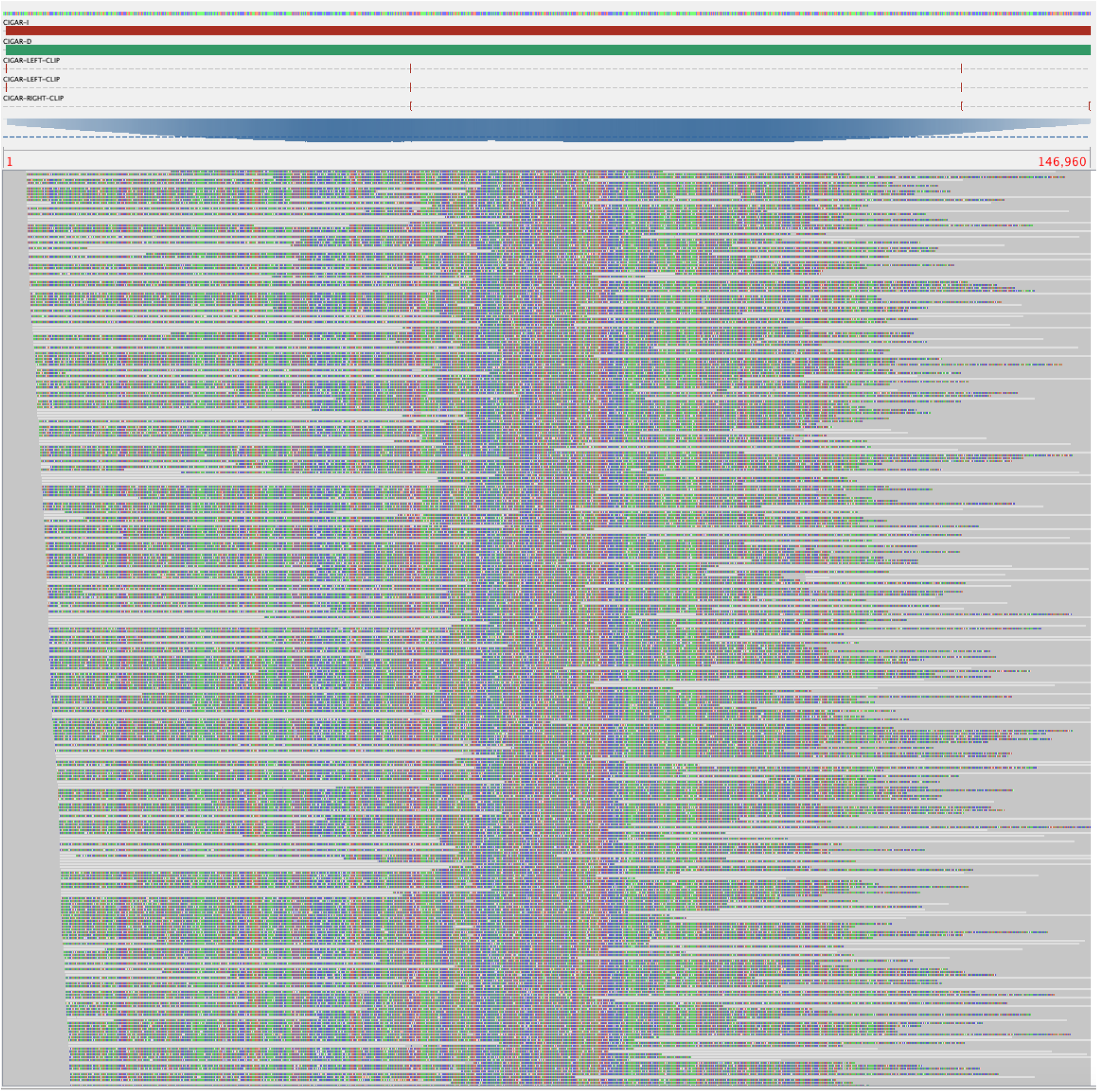
Genomic alignments of Pacbio long reads to the mitochondrial scaffold generated via Canu assembly in order to confirm the length of the circular mitogenome of the *C. posadasii*, strain Silveira.

**Suppl Figure 3:**
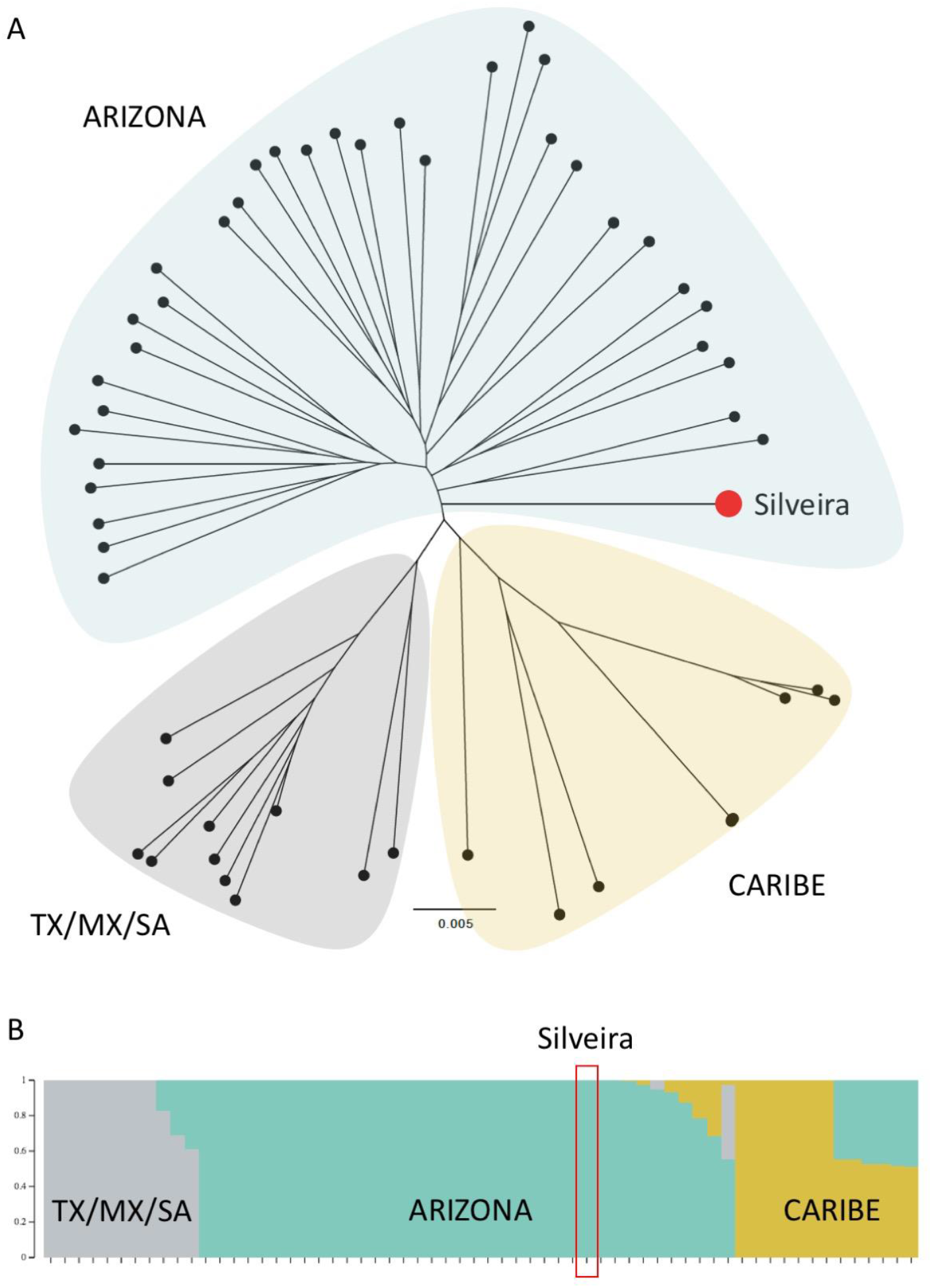
Whole-genome genotyping of the Silveira strain. A) Genome-level tree of *C. posadasii* based on Maximum likelihood (ML) method. The tree is unrooted and phylogenetic clades are displayed proportionally to the branch length. Dual branch support was evaluated using 1,000 ultrafast bootstraps coupled with a Shimodaira–Hasegawa-like approximate likelihood ratio test and were used to infer the observed phylogenetic partition. The phylogenetic groups were defined and are displayed along the tree. B) Structure plots of *C. posadasii* showing the presence of three well defined populations as follows: ARIZONA, CARIBE and TX/MX/SA. Each strains is represented by a single vertical bar, which is partitioned into K=3 and the Silveira strain is nested within the ARIZONA population. A single color represents an absence of admixture, which is the case for the Silveira strain that is highlighted in red.

